# Deleterious variants in Asian rice and the potential cost of domestication

**DOI:** 10.1101/057224

**Authors:** Qingpo Liu, Yongfeng Zhou, Peter L. Morrell, Brandon S. Gaut

## Abstract

Many SNPs are predicted to encode deleterious amino acid variants. These slightly deleterious mutations can provide unique insights into population history, the dynamics of selection, and the genetic bases of phenotypes. This is especially true for domesticated species, where a history of bottlenecks and selection may affect the frequency of deleterious variants and signal a ‘cost of domestication’. Here we investigated the numbers and frequencies of deleterious variants in Asian rice (*O. sativa*), focusing on two varieties (*japonica* and *indica)* and their wild relative (*O. rufipogon*). We investigated three signals of a potential cost of domestication in Asian rice relative to *O. rufipogon*: an increase in the frequency of deleterious SNPs (dSNPs), an enrichment of dSNPs compared to synonymous SNPs (sSNPs), and an increased number of deleterious variants. We found evidence for all three signals, and domesticated individuals contained ~3-4% more deleterious alleles than wild individuals. Deleterious variants were enriched within low recombination regions of the genome and experienced frequency increases similar to sSNPs within regions of putative selective sweeps. A characteristic feature of rice domestication was a shift in mating system from outcrossing to predominantly selfing. Forward simulations suggest that this shift in mating system may have been the dominant factor in shaping both deleterious and neutral diversity in rice.

## Introduction

Several studies have suggested that there is a “cost of domestication” (Schubert et al., 2014), because crops may harbor slightly deleterious mutations that reduce their relative fitness (Lu et al., 2006). Under this hypothesis, the decreased effective population size (N_e_) during a domestication bottleneck reduces the efficacy of genome-wide selection (Charlesworth and Willis, 2009), leading to an increase in the frequency of slightly deleterious variants (Lohmueller et al., 2008, Casals et al., 2013). The fate of these variants also relies on linkage, because selection is less effective in genomic regions of low recombination (Hill and Robertson, 1966, Felsenstein and Yokoyama, 1976) and because deleterious variants may hitchhike with alleles that are positively selected for agronomic traits (Fay and Wu, 2000, Hartfield and Otto, 2011, Campos et al., 2014). Overall, the cost of domestication is expected to increase the frequency of deleterious variants in small relative to large populations, in regions of low recombination, and near sites of positive selection.

This hypothesis about the cost of domestication parallels the debate regarding the genetic effects of migration-related bottlenecks and demographic expansion in human populations (Lohmueller et al., 2008, Casals et al., 2013, Peischl et al., 2013, Simons et al., 2014). The debate regarding human populations is contentious, perhaps because it suggests that some human populations may, on average, carry a greater load of deleterious variants than others (Peischl et al., 2016). Studies in humans also suggest that subtlety of interpretation is required when considering the relative frequency of deleterious variants in populations, because both the effect size and the relative dominance of deleterious variants likely play a role in how mutations impact the fitness of populations (Henn et al., 2016). Moreover, deleterious variants in non-equilibrium populations, such as those that have experienced a recent bottleneck, may return to pre-bottleneck frequencies more rapidly than neutral variants (Brandvain and Wright, 2016). It nonetheless remains an important task to identify the number, frequency and genomic distribution of deleterious variants in humans, for the purposes of disentangling evolutionary history and for understanding the association between deleterious variants and disease (Kryukov et al., 2007, Eyre-Walker, 2010, Gazave et al., 2013, Lohmueller, 2014a, Simons et al., 2014, Uricchio et al., 2016).

In plant crops, the potential for a ‘cost of domestication’ was first examined in Asian rice (*O. sativa*) (Lu et al., 2006). At the time, limited population resequencing data were available, so Lu et al (2006) compared two *O. sativa* reference genomes to that of a related wild species (*O. brachyntha*). They found that the *K_a_*/*K_s_* ratio for radical, presumably deleterious amino acid variants was higher between the two *O. sativa* genomes than between *O. sativa* and *O. brachyntha*. The *K_a_*/*K_s_* ratios for individual genes were negatively correlated with genomic recombination rates, potentially suggesting hitchhiking effects (Lu et al., 2006). Finally, they showed that deleterious amino acid variants in rice were typically found at intermediate population frequencies. Altogether, they hypothesized that these observations reflect a cost of domestication, whereby deleterious variants are enriched (relative to synonymous variants) during domestication. They hypothesized that enrichment was driven by two evolutionary processes: relaxation of selective constraint and hitchhiking due to artificial selection.

A handful of studies have since analyzed deleterious variants in crops based on resequencing data (Gunther and Schmid, 2010, Nabholz et al., 2014, Renaut and Rieseberg, 2015, Kono et al., 2016). Together these studies report that: i) deleterious variants are found at higher population frequencies within crops compared to their wild relatives and ii) the relative frequency of deleterious to neutral variants is higher in crops than in their wild progenitors. For example, Renaut and Rieseberg (2015) measured the proportion of deleterious SNPs to synonymous SNPs in wild and cultivated accessions of sunflower, and they showed that this proportion was consistently higher for domesticated than for wild accessions. More limited analyses have also shown that deleterious variants are enriched within genes associated with phenotypic traits (Mezmouk and Ross-Ibarra, 2014, Kono et al., 2016), suggesting both that deleterious variants are affected by selection through hitchhiking and that the study of deleterious variants is crucial for understanding the potential for crop improvement (Morrell et al., 2011). While a general picture is thus beginning to emerge, most of these studies have suffered from substantial shortcomings, such as small numbers of genes, low numbers of individuals, or the lack of an outgroup to infer ancestral states. Moreover, no study of crops has yet investigated the frequency of deleterious variants in putative selective sweep regions, which is especially important given the hypothesis that artificial selection has increased the frequency of deleterious mutations (Lu et al., 2006).

In this study, we reanalyze genomic data from hundreds of accessions of Asian rice and its wild relative *O. rufipogon*. Asian rice feeds more than half of the global population (IRGSP, 2005), but the domestication of the two main varieties of Asian rice (ssp. *japonica* and ssp. *indica*) remains enigmatic. It is unclear whether the two varieties represent independent domestication events (Londo et al., 2006, Civian et al., 2015), a single domestication event with subsequent divergence (Gao and Innan, 2008, Molina et al., 2011), or separate events coupled with substantial homogenizing gene flow of beneficial domestication alleles (Caicedo et al., 2007, Sang and Ge, 2007, Zhang et al., 2009, Huang et al., 2012a, b). It is clear, however, that domestication has included a shift in mating system from predominantly outcrossing *O. rufipogon* [which has outcrossing rates between 5% and 60%, depending on the population of origin and other factors (Oka and Miroshima, 1967)] to predominantly selfing rice [which has outcrossing rates of ~1% (Oka, 1988)]. This shift in mating system has the potential to affect the population dynamics of deleterious variants, because inbreeding exposes partially recessive variants to selection (Lande and Schemske, 1985), which may in turn facilitate purging of deleterious alleles (Arunkumar et al., 2015).

Commensurate with its agricultural importance, the population genetics of Asian rice have been studied in great detail. Resequencing studies indicate that nucleotide sequence diversity is ~2 to 3-fold lower in *indica* rice compared to *O. rufipogon* (Caicedo et al., 2007, Huang et al., 2012b) and that diversity in *indica* is ~2 to 3-fold higher than *japonica* rice (Zhu et al., 2007, Huang et al., 2012b). *Japonica* rice is often further separated into tropical and temperature germplasm, with higher diversity in the former (Caicedo et al., 2007). Sequence polymorphism data have also shown that the derived site frequency spectrum (SFS) of rice varieties exhibit a distinct U-shaped distribution relative to *O. rufipogon,* due either to the genome-wide effects of selection or migration (Caicedo et al., 2007). However, the population genetics of putatively deleterious variants have not been studied across *O. sativa* genomes, nor have deleterious variants been contrasted between *O. sativa* and *O. rufipogon* based on genomic data.

Here we assess whether genomic data provide evidence for a “cost of domestication” in rice. We consider three measures of cost, as defined previously in the literature. The first is elevated population *frequencies* of deleterious variants that remain after domestication (Lu et al., 2006); the second is an enrichment in the *proportion* of deleterious SNPs to synonymous SNPs in cultivated vs. wild individuals (Renaut and Rieseberg, 2015); and the third is an increase in the *number* of derived deleterious variants in domesticated vs. wild germplasm. To our knowledge, this last measure of cost has not yet been considered in the context of crop domestication. We include it here because it is central to discussions of deleterious mutations in human populations, particularly with regard to population expansion (e.g., Lohmueller, 2014b, Simons et al., 2014, Henn et al., 2016).

To identify putatively deleterious variants, we have utilized two *O. sativa* datasets: one with many accessions (*n* = 766) but low sequencing coverage (1-2X), and the other with fewer individuals (*n* = 45) but enhanced coverage. For both datasets, we re-map raw reads and then apply independent computational pipelines for SNP variant detection. We have used two different approaches – PROVEAN (Choi et al., 2012) and SIFT (Kumar et al., 2009) - to predict which nonsynonymous SNPs are deleterious. With these predicted deleterious variants, we investigate three signals of cost (i.e., frequencies, enrichment and numbers of deleterious variants). We also examine the distribution of deleterious variants relative to genome-wide recombination rates and the locations of putative selective sweeps. Finally, we attempt to gain insights into the relative contributions of demography, linkage, positive selection and inbreeding on the dynamics of deleterious variants within Asian rice.

## Results

### Datasets and site frequency spectra

To investigate the population dynamics of deleterious variants, we collated two rice datasets. The first was based on the genomic data of 1,212 accessions reported in Huang et al. (2012b) (Table S1). This dataset, which we call the ‘BH’ data after the senior author, contained raw reads from 766 individuals of Asian rice, including 436 *indica* accessions and 330 *japonica* accessions. The BH dataset also included 446 accessions representing three populations of *O. rufipogon*, the wild ancestor of cultivated rice (Table S1). Huang et al. (2012b) determined that their *O rufipogon* accessions represented three different wild populations, which we denote W_I_, W_II_ and W_III_. They also inferred that W_I_ was ancestral to *indica* rice and that W_III_ was ancestral to *japonica* rice. Accordingly, we based our cultivated-to-wild comparisons on *indica* vs. W_I_ and *japonica* vs. W_III_ for the BH data, but when appropriate we also included comparisons to the complete set of wild accessions (W*all*). For these BH data, we remapped sequencing reads to the *japonica* reference sequence (Goff et al., 2002), then used ANGSD (Korneliussen et al., 2014) to apply cut-offs for quality and coverage and to estimate the SFS (see Materials and Methods).

The second dataset, which we call the ‘3K’ data (Li et al., 2014), consisted of 15 cultivated, high-coverage (>12X) accessions for each of *indica*, tropical *japonica*, and temperate *japonica* (Table S2). We also included data from the BH dataset of the 15 wild *O. rufipogon* individuals with the highest coverage, which we denote W_15_; coverage for the W_15_ individuals ranged from 4.6X to 9.8X. For this dataset, reads were again mapped to the *japonica* reference, but SNPs were called using tools from GATK and SAMtools (see Materials and Methods). For many analyses, we focused on a subset of this 3K data (3K_subset_) that included only sites without missing data and for which SNPs were identified within the entire *n*=60 sample, rather than within individual taxa.

Once identified, we annotated SNPs as either non-coding (ncSNPs), synonymous (sSNPs), Loss of Function (LoF) or nonsynonymous. LoF SNPs were those that contribute to apparent splicing variation, the gain of a stop codon or the loss of a stop codon. Nonsynonymous SNPs were predicted to be tolerant (tSNPs) or deleterious (dSNPs) based on PROVEAN (Choi et al., 2012) or SIFT (Ng and Henikoff, 2003). Table S3 reports raw numbers of detected SNPs in both datasets. In the BH rice samples, we identified hundreds of LoF mutations and predicted 7,506 and 4,530 dSNPs in *indica* and *japonica* samples using PROVEAN. Despite fewer accessions, we identified more SNPs within the 3K data owing to higher sequence coverage, including 21,234 dSNPs in *indica* rice (Table S3).

To determine the unfolded site frequency spectra for various datasets and SNP classes, we defined SNPs as ancestral or derived based on comparison to 93 *O. barthii* accessions (Table S4). For the BH data, we reduced the sample size to 70 for each population, based on sampling and coverage criteria (Materials and Methods). The resulting SFS had a U-shape for all SNP categories in cultivated rice, as observed previously (Caicedo et al., 2007), but not for ancestral *O. rufipogon* (Figures 1 & S1). The SFS differed significantly between wild and domesticated samples for all SNP categories (Kolmogorov-Smirnoff tests; p<0.001; Figures 1 & S1).

**Figure 1:**
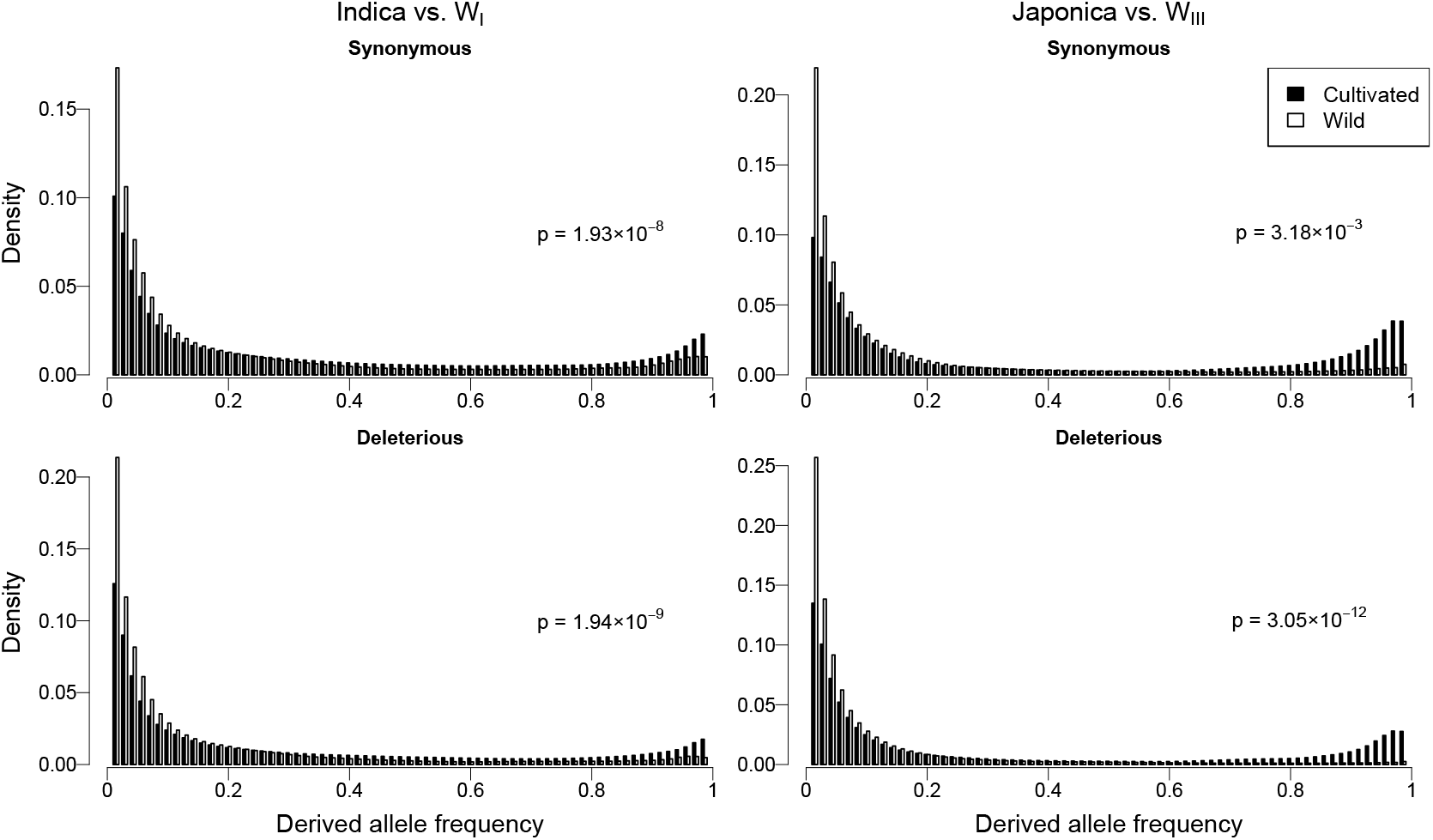
The site frequency spectrum (SFS) for cultivated rice and *O. rufipogon*, based on BH data. The top row represents sSNPs, and the bottom row represents dSNPs. Additional SNP classes are graphed in Figure S1. The two columns represent *indica* rice on the left and *japonica* rice on the right. As per Huang et al (2012b), *indica* rice is contrasted to the accessions from wild population I (WI) and *japonica* rice is contrasted to wild sample population III (W_III_). The Density on the y-axis is the proportion of alleles in a given allele frequency. Each graph reports the *p*-value of the contrast in SFS between cultivated and wild samples.

SNPs in the BH data were based on detecting polymorphisms within each taxon separately (Table S3), which limits the potential to infer sites at the extremes of the SFS – i.e, the zero and fixed classes. To estimate these classes, we focused on the 3K_subset_ data, which had 2,239,824 SNPs across the 60 individuals, including 22,377 dSNPs, 65,594 tSNPs, 81,648 sSNPs and 4,102 LoF variants (see also Table S3). Nucleotide diversity estimates (π) for noncoding and 4-fold degenerate sites based on these data were similar to those of previous studies (Huang et al., 2012b), with the expected hierarchy of diversity relationships (i.e., *O. rufipogon > indica > tropical japonica > temperate japonica*) (Caicedo et al., 2007) (Table 1). With the 3K_subset_ data, the comparisons between the wild and cultivated SFS were not significant (Kolmogorov-Smirnoff; p>0.05), but the resulting SFS were similar to the BH data in exhibiting hints of a U-shaped SFS for all three cultivated taxa and for most site categories (Figures 2 & S2). This U-shape included enhanced frequencies of fixed and high frequency (>12) derived variants and a dearth of low frequency (<3) variants in domesticates compared to the W_15_ sample (Figures 2 & S2). These comparisons also illustrate that the zero class was greatly enhanced in domesticated taxa, indicative of the loss of rare, low frequency variants.

**Figure 2:**
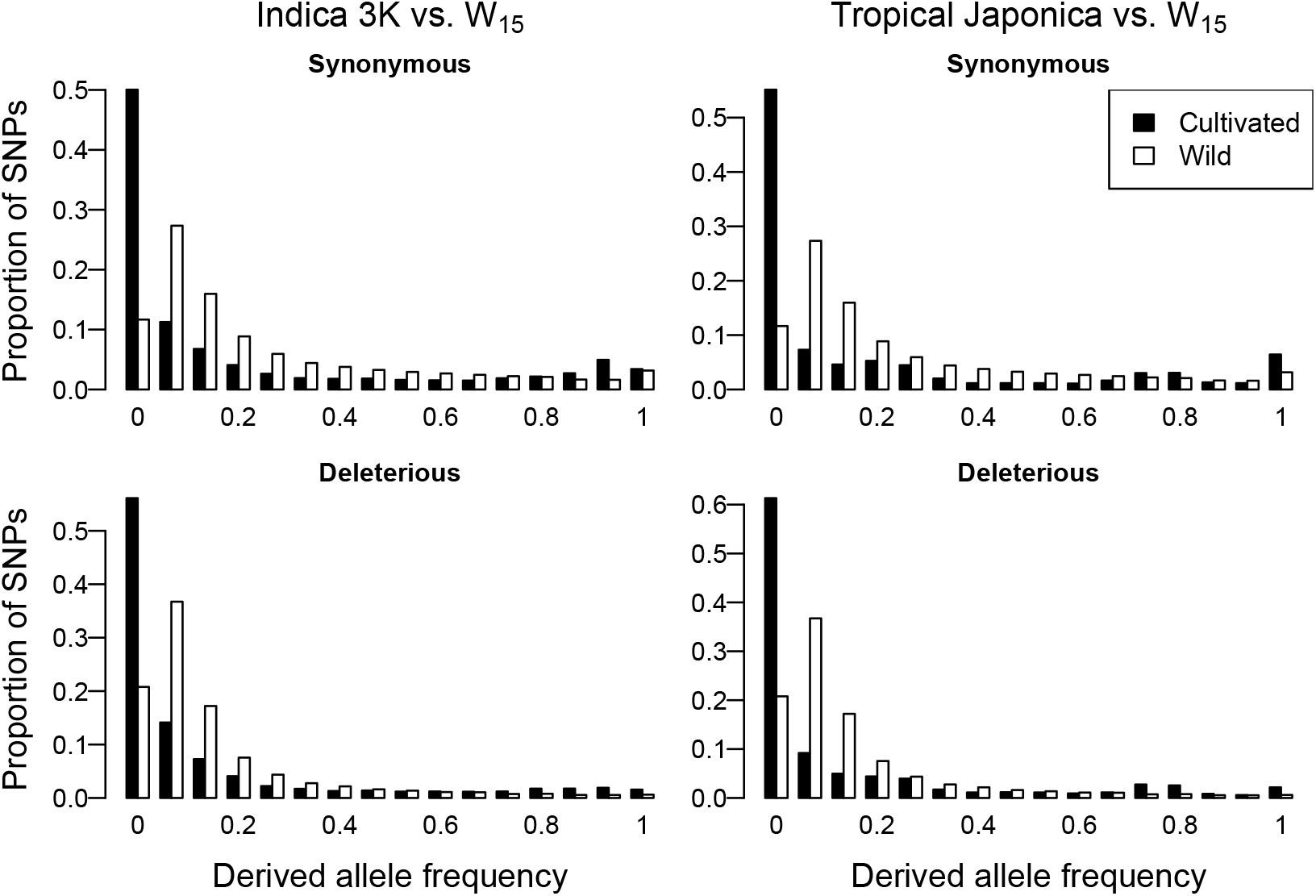
The SFS for cultivated rice and *O. rufipogon*, based on 3K_subset_ data for the *indica* and tropical *japonica* samples. The top row represents sSNPs, and the bottom row represents dSNPs. Additional SNP classes are graphed in Figure S2. The two columns represent *indica* rice on the left and tropical *japonica* rice on the right; temperate *japonica* is included in Figure S2.

**Table 1:** Comparison of genetic diversity (π), average counts of derived deleterious variants (#dSNPs) per individual, and the average ratio of deleterious to synonymous variants (#dSNPs/#sSNPs) per individual.

| Sample <sup>a</sup> | $\pi_{\text{noncoding}}$ <sup>b</sup> | $\pi_{4\text{-fold}}$ <sup>b</sup> | Avg.<br>#dSNPs <sup>c</sup> | Avg.<br>[#dSNPs /#sSNPs] <sup>d</sup> |
| --- | --- | --- | --- | --- |
| *indica* | 0.00394 | 0.00289 | 6351.4<br>($\pm 76.3$) | 0.1818<br>($\pm 0.0032$) |
| *japonica*<br>(*temperate*) | 0.00252 | 0.00177 | 6290.3<br>($\pm 59.7$) | 0.1912<br>($\pm 0.0020$) |
| *japonica*<br>(*tropical*) | 0.00345 | 0.00246 | 6300.7<br>($\pm 36.9$) | 0.1880<br>($\pm 0.0031$) |
| W<sub>15</sub> | 0.00620 | 0.00505 | 6098.5<br>($\pm 126.8$) | 0.1522<br>($\pm 0.0067$) |

<sup>a</sup> All samples are based on the 3K<sub>subset</sub> data.
<sup>b</sup> Pairwise diversity, based on a common set of SNPs without missing data, along with invariant sites without missing data.
<sup>c</sup> The number of derived segregating and fixed dSNPs per individual, averaged over the 15 individuals within each of the four groups. Parentheses indicate the 95% confidence interval.
<sup>d</sup> The average ratio of derived dSNPs to sSNPs per individual. Parentheses indicate the 95% confidence interval.

Inferred shifts in the SFS after domestication were robust to: *i*) dataset, because the 3K_subset_ and BH datasets yielded similar results, *ii*) SNP calling approaches, because different methods were applied to the 3K and BH datasets, *iii*) the composition of the wild sample, because similar patterns were observed when the BH *japonica* and *indica* samples were compared to W_*all*_ (*p* ≤ 1.93 × 10^-8^ for all comparisons in both varieties) (Figure S3), *iv*) variation in sample sizes (*n*) among taxa, because the BH data did not have the same number of individuals per taxon, while the 3K data did (Table S3), and *v*) the prediction approach used to identify dSNPs (i.e., PROVEAN or SIFT; Figures S4 & S5). Overall, the derived variants that remained after domestication were shifted to higher frequency, as is expected following a bottleneck (Simons et al., 2014).

### Enhanced frequencies and numbers of deleterious variants in rice

Previous work has defined the cost of domestication as higher frequencies of dSNPs (Lu et al., 2006), particularly differential shifts in frequencies of dSNPs relative to putatively neutral SNPs (Gunther and Schmid, 2010). To investigate frequency shifts, we plotted the ratio of the number of derived dSNPs vs. derived sSNPs for each frequency category of the SFS. Figure 3 shows that, for the BH data, both *indica* and *japonica* have enhanced numbers of derived dSNPs to sSNPs relative to *O. rufipogon* across the entire frequency range (Wilcoxon rank sum: *indica* vs. W_I_, *p* = 4.98 × 10^-16^; *japonica* vs. W_III_, *p* < 2.20 × 10^-16^; Figure 3). The 3K_subset_ data exhibited similar properties throughout most of the frequency range, with the exception of the lowest frequency classes, but the distributions remained significantly different overall (Wilcoxon rank sum: *indica* 3K vs. W_15_, *p* = 0.035; tropical *japonica* 3K vs. W_15_, *p* = 0.023; Figure 3).

**Figure 3:**
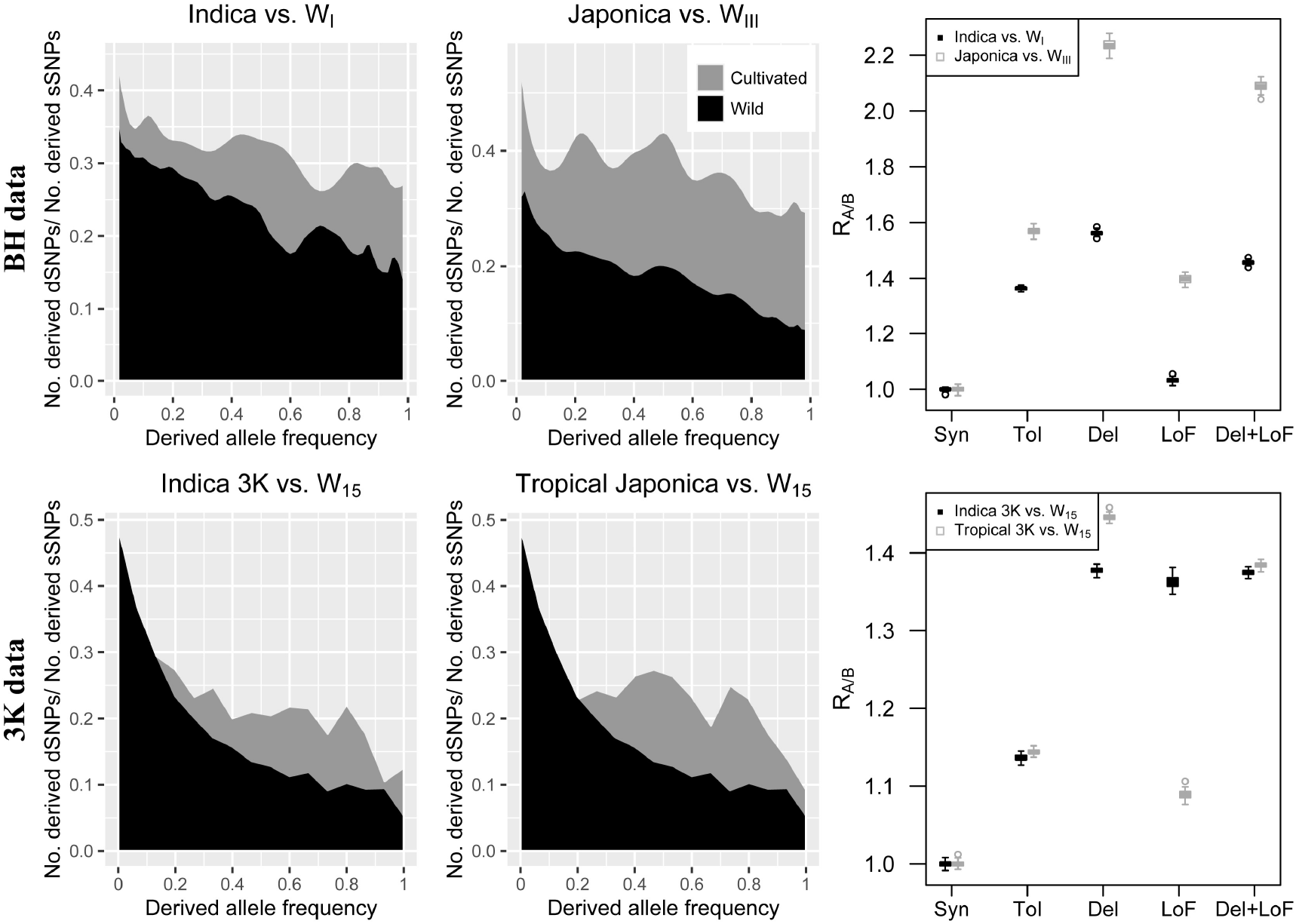
Comparisons of the number of derived dSNP to sSNP between wild and cultivated samples based on their frequencies. The top row reports results based on the BH data. From left to right, the panels represent: *left*) the ratio of the number of dSNPs to sSNPs (*y*-axis) at each derived allele frequency (*x*-axis) between *indica* rice and the W_I_ sample; *middle*) the ratio of the number of dSNPs to sSNPs (y-axis) at each derived allele frequency (*x*-axis) between *japonica* rice and the W_III_ sample and *right*) a measure R_(A/B)_ of the relative accumulation of SNPs in *indica* or *japonica* rice compared to *O. rufipogon*, where a value > 1.0 indicates an increased population density of that SNP type relative to wild rice. Bars indicate standard errors. The bottom row reports the 3K_subset_ data, and the three panels (left to right) are equivalent to those from the BH data.

We also calculated R_(A/B)_, a measure that compares the frequency and abundance of dSNPs vs. sSNPs in one population (A) relative to another (B) (Xue et al., 2015). When R_(A/B)_ is > 1.0, it reflects an overabundance of derived dSNPs (or LoF variants) relative to sSNPs in one population over another across the entire frequency range. As expected from SFS analyses, we found that R_(A/B)_ was > 1.0 for LoF variants and for dSNPs in *indica* relative to the W_I_ population (*p* ≤ 2.30 × 10^-139^ for all three comparisons; Figure 3) and in *japonica* relative to W_III_ (*p* ~ 0.000 for the three comparisons; Figure 3). The 3K_subset_, which included both the zero and fixed classes of variants, yielded similar results (*p* ~ 0.000 for all six comparisons; Figure 3). Hence, all cultivated samples contained increased proportions of derived dSNPs to derived sSNPs relative to wild samples.

An enrichment of the number of derived dSNPs to sSNPs within cultivated individuals has also been cited as evidence for a cost of domestication (Renaut and Rieseberg, 2015). Because the 3K_subset_ included the same sample size and number of nucleotide sites across taxa, it permitted direct comparisons of the numbers of derived alleles across individuals. We therefore calculated the average number of derived deleterious variants within individuals and the average ratio of derived dSNPs to sSNPs within individuals (Table 1). Following Henn et al. (2016), we counted the number of derived variants per individual as the number of heterozygous sites plus twice the number of derived homozygous sites. We included both fixed and segregating derived variants in our calculations.

The results showed that the count of derived, putatively deleterious variants *increased* for each cultivated individual, on average, despite lower overall non-coding and synonymous diversity (π) in the cultivated samples (Table 1). These counts differed significantly between the W_15_ sample and the three cultivated samples for dSNPs (Mann-Whitney *U*, *p* ≤ 0.0344), representing a ~3 to 4% increase in the average number of derived deleterious variants per individuals. Accordingly, the ratio of derived deleterious to synonymous variants per individual was significantly higher for the domesticated rice samples than for *O. rufipogon* (Mann-Whitney *U*, *p* ≤ 2.51 × 10^-6^ for all three comparisons). Analysis based on SIFT prediction of deleterious variants yielded similar trends (Table S5).

### dSNPs are enriched in regions of low recombination

We have established that our rice samples have enhanced frequencies, proportions and numbers of dSNPs. We now evaluate whether the accumulation of putatively deleterious variants was homogenous across the genome. Theory predicts that diversity should be lower in low recombination regions (Begun and Aquadro, 1992, Charlesworth, 1994) and also that fate of dSNPs relative to sSNPs may differ between high and low recombination regions due to interference (Felsenstein, 1974b). To test these predictions, we used a genetic map to calculate recombination rate in windows across rice chromosomes. Applying 2MB windows to the 3K data, we found that the average number of derived (segregating + fixed) sSNPs and dSNPs per individual were significantly positively correlated with recombination rate in each of the rice samples (Figure 4; *p*≤5.18 × 10^-8^), indicating reduced diversity in low recombination regions.

**Figure 4:**
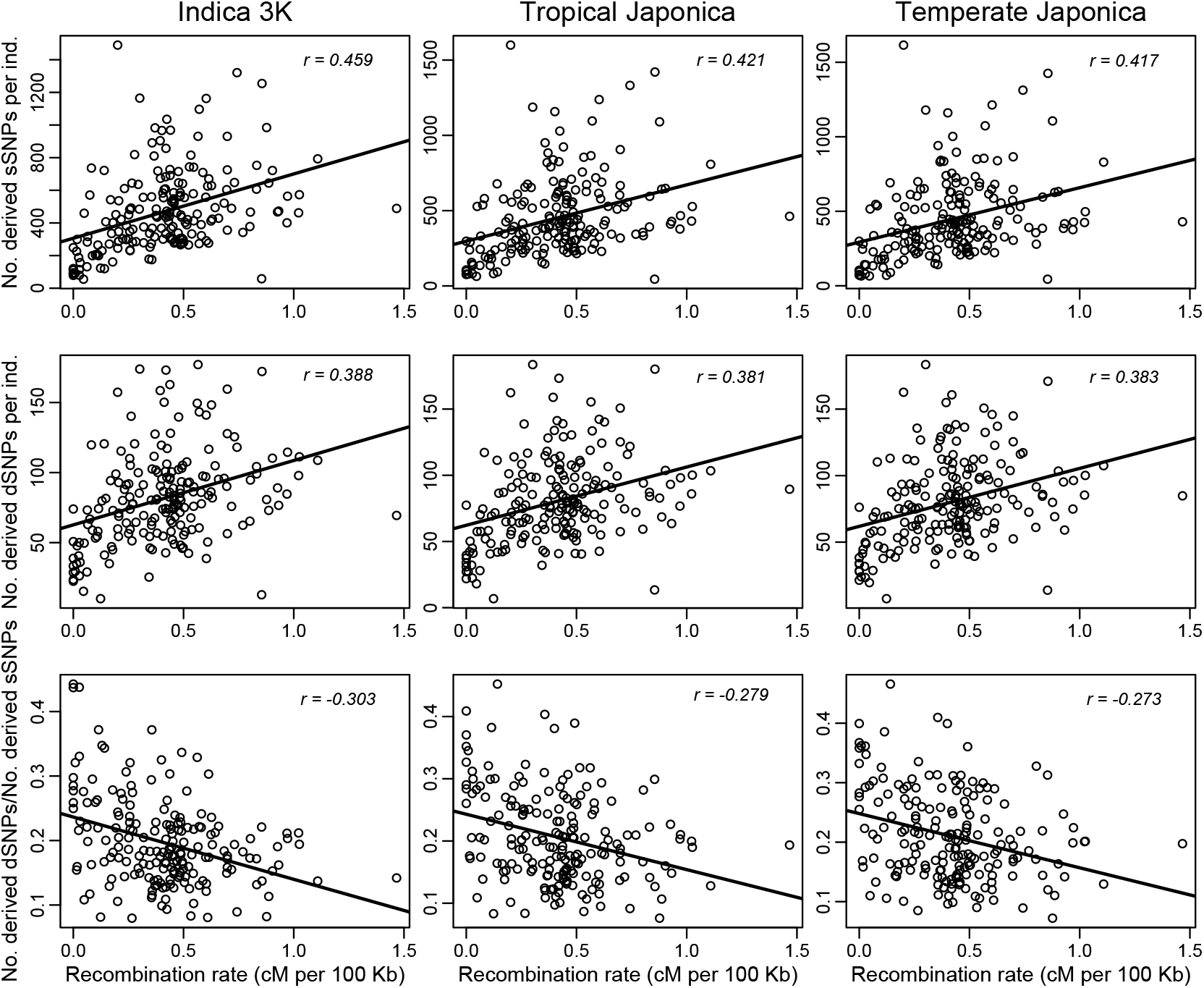
Patterns of genomic variation relative to recombination, based on the 3K data. The x-axis for each graph is the recombination rate (x-axis) as measured by centiMorgans (cM) per 100 kb. The y-axis varies by row. The top row is the average number of derived (segregating + fixed) synonymous variants per individual, as measured by in 2MB windows; the middle row is the average number of derived (segregating + fixed) deleterious variants in 2MB windows; and the bottom row is the ratio of derived dSNPs to sSNPs in 2MB windows. The *p*-values for Spearman correlations for the top row of graphs are all *p* ≤ 2.1 × 10^-9^; the *p*-values for the middle row are all *p* ≤ 5.2 × 10^-8^; and the *p*-values for the bottom row are all *p* ≤ 1.3 × 10^-4^. A table for similar analyses with the full BH and 3K dataset are provided in Tables S6 and S7.

Following precedent (Renaut and Rieseberg, 2015, Kono et al., 2016), we also plotted recombination rate against the ratio of the number of derived dSNPs to derived sSNPs; these correlations were significantly negative (*p*≤1.32 × 10^-4^). For completeness, we repeated these analyses on the density and pairwise diversity of variants in the full BH and 3K datasets, which yielded similar results (Tables S6 and S7). In short, the ratio of derived dSNPs to derived sSNPs was consistently higher in regions of low recombination.

### dSNP frequencies in regions of putative selective sweeps

Regions linked to selective sweeps (SS) may have increased frequencies of derived mutations (Fay and Wu, 2000), including dSNPs (Hartfield and Otto, 2011). Consistent with this expectation, a previous study of domesticated dogs has shown that the frequency of both dSNPs and sSNPs are inflated within SS regions (Marsden et al., 2016). Prompted by these observations, we investigated the distribution of deleterious and synonymous variants in putative SS regions, to test two hypotheses. The first was that SS regions have increased frequencies of derived SNPs relative to the remainder of the genome. The second was that SS regions alone explain the accumulation of high frequency derived dSNPs in Asian rice.

We used three different approaches to identify SS regions. First, we used the SS regions defined by Huang et al. (2012c), which were based on the relative difference in π between wild and domesticated populations (Huang et al., 2012c). That is, the regions were based on π_*d*_/π_*w*_, where π is measured per base pair, and the subscripts refer to domesticated and wild samples. We also inferred selective sweeps using two additional approaches: SweeD (Pavlidis et al., 2013) and XP-CLR (Chen et al., 2010). SweeD identifies regions of skewed SFS relative to background levels for a single population (i.e., the rice sample). In contrast, XP-CLR searches for genomic regions for which the change in allele frequency between two populations (cultivated vs. wild samples) occurred too quickly at a locus, relative to the size of the region, to be caused by genetic drift. Both SweeD and XP-CLR were applied with a 5% cutoff. Because XP-CLR requires explicit genotypes, we used the 3K datasets for all of the SS analyses (Methods).

Focusing on the *indica* 3K dataset for simplicity, the three approaches identified different numbers, locations and sizes of selective sweeps (Table 2). For example, Huang et al. (2012c) defined 84 SS regions that encompassed 9.98% of the genome. In contrast, SweeD identified 485 SS regions, and XP-CLR distinguished an intermediate number of 161 SS regions. Consistent with the 5% cutoff, SweeD and XP-CLR identified 4.61% and 5.02% of the genome, respectively, as having been under selection (Table 2). To see if the same genes were identified with different SS identification methods, we calculated the degree of overlap across methods, focusing on the percentage of genes that two methods identified in common (see Methods). The overlap was surprisingly low (Figure 5 & S6-S16). Across the entire genome, the putative SS regions defined by SweeD and Huang et al (2012c) shared 6.24% of genes. Similarly, the regions defined by XP-CLR shared 8.51% and 8.69% of genes with Huang et al. (2012c) and SweeD, respectively.

**Figure 5:**
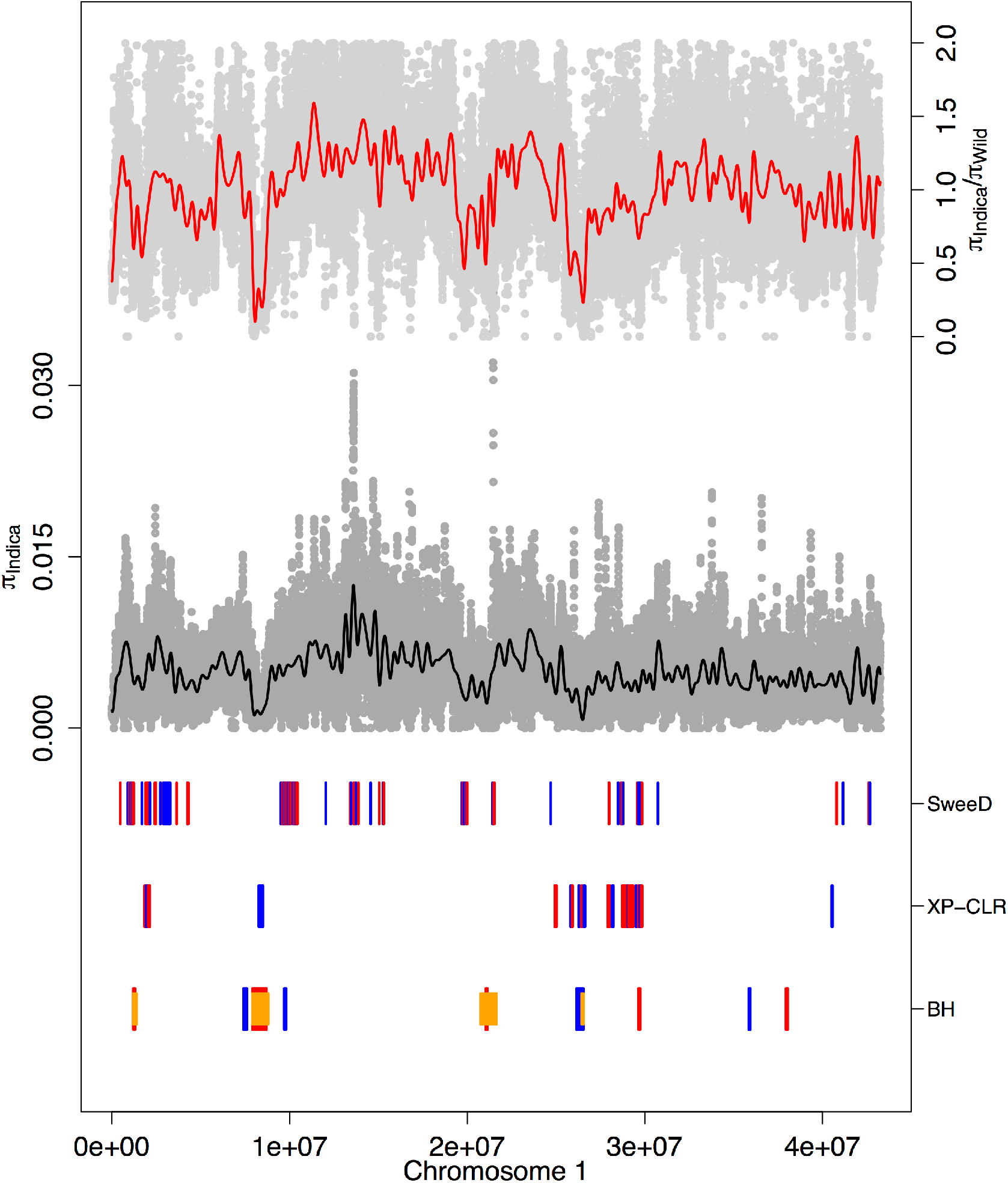
A graph of the location of inferred SS regions along Chromosome 1 for the 3K *indica* dataset. The x-axis is the location along the chromosome, measured in base pairs. The top graph (red) indicates the ratio of π for the *indica* accessions against a set of wild accessions. The (grey) background represents values for windows of 10kb with a step size of 1kb. Values > 2.0 were omitted for ease of presentation, and the line was smoothed. The middle graph shows values of π for the *indica* accessions. The bars at the bottom represent inferred SS regions using SweeD and XP-CLR, along with predefined SS regions (BH) defined by Huang et al. (2012b). The red and blue colors are included to help differentiate SS regions; the orange bars represent additional SS regions defined by Huang et al. (2012b) on the basis of their combined *indica+japonica* dataset. The width of each bar is proportional to the length of the corresponding SS region along chromosome. Similar graphs for chromosomes 2 through 12 are available as supplemental figures (Figures S6 to S16).

**Table 2:** The number and percentage of SS regions identified by different methods, based on 3K data.

|  | *indica* |  | *japonica*<br>(temperate) |  | *japonica* (tropical) |  |
| --- | --- | --- | --- | --- | --- | --- |
|  | No. | Extent <sup>b</sup> | No. | Extent <sup>b</sup> | No. | Extent <sup>b</sup> |
| Huang et al<br>(2012b) | 84 <sup>a</sup> | 9.98% | 103 <sup>c</sup> | 15.32% | 103 <sup>c</sup> | 15.32% |
| SweeD | 485 | 4.61% | 461 | 4.76% | 389 | 4.81% |
| XP-CLR | 161 | 5.02% | 160 | 8.41% | 171 | 5.62% |

<sup>a</sup> Based on 60 SS regions identified as specific to *indica*, which overlapped with 31 of 55 regions identified in the combined samples of *indica* and *japonica* rice, for a total of $[60+(55-31)]=84$ .
<sup>b</sup> Extent = the percentage of the reference genome covered by SS regions.
<sup>c</sup> Based on 62 SS regions identified as specific to *japonica*, which overlapped with 14 of 55 regions identified in the combined samples of *indica* and *japonica* rice, for a total of $[62+(55-14)]=103$ .

To determine if SS regions have increased frequencies of derived dSNPs, we contrasted the SFS between SS and non-SS regions for derived segregating and fixed sSNPs and dSNPs (see Marsden et al., 2016). The SFS were skewed for SS regions relative to non-SS regions for both SNP classes, independent of the method used to detect selective sweeps (Figure 6A). We summarized the shift in frequencies by counting the number of derived alleles (DAC) per SNP (Figure 6B) (Marsden et al., 2016), which showed that SS regions also contained higher DACs (Figure 6B). Note that these results were not completely unexpected, because the methods used to define SS regions rely, in part, on identifying a skewed SFS relative to the genomic background (see Discussion).

**Figure 6:**
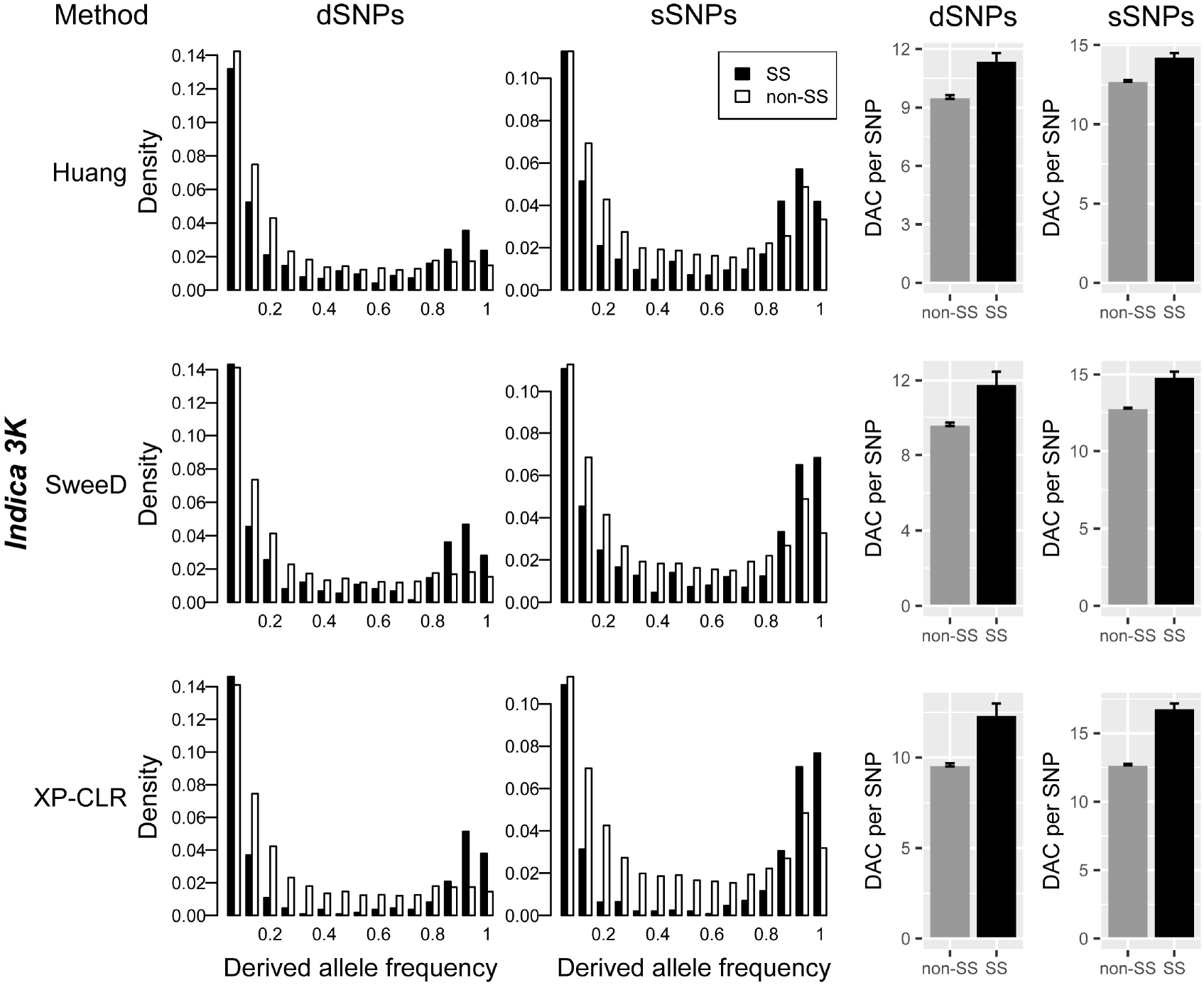
A comparison between selective sweep (SS) and non-SS regions based on the *indica* 3K_subset_ dataset. The rows correspond to different methods employed to detect sweeps, including SS regions from Huang et al (2012b) (top row), SweeD (middle row), and XP-CLR (bottom row). The set of histograms on the right compare the derived allele count (DAC) of segregating and fixed synonymous site or putatively deleterious sites between SS regions and the remainder of the genome (non-SS regions).

Did sweeps affect dSNPs more or less than sSNPs? To investigate this question, we calculated the ratio of the mean DAC for SS and non-SS regions. There was some variation among SS methods. For example, the SS regions exhibited a 1.23-fold enrichment for dSNPs vs. a slightly smaller 1.16-fold enrichment for sSNPs when SS regions were based on SweeD (Table S8). Similarly, the SS regions defined by Huang et al. (2012c) included a 1.20- and 1.12-fold enrichment for dSNPs and sSNPs, respectively. SS regions defined by XP-CLR showed the reverse: slightly higher enrichment for sSNPs (1.33) than for dSNPs (1.29). Altogether, the extent to which hitchhiking drove dSNPs and sSNPs to higher frequency seems to be roughly equivalent.

Enhanced SNP frequencies in SS regions raise the possibility that selective sweeps alone explain the shifted SFS of *indica* rice relative to *O. rufipogon*. To examine this second hypothesis, we removed all SS regions (as defined by SweeD, XP-CLR and π_*d*_/π_*w*_) from the *indica* 3K dataset and recalculated the SFS. Even with SS regions removed, the SFS for wild and cultivated samples remained significantly different for sSNPs and dSNPs (*p* ≤ 0.0067). These results imply either that positive selection is not the only cause of the U-shaped SFS in *indica* rice (Caicedo et al., 2007) or that linked selection has affected more of the genome than is encompassed within the identified SS regions.

We have reported results based on *indica* rice, but we also performed analyses of SS regions for the 3K temperate and tropical *japonica* datasets (Table 2). The results were similar to *indica* rice in three respects. First, although a greater extent of the genome tended to be identified as SS regions in *japonica* (Table 2), the overlap among SS regions identified by different methods was again low (< 9%). Second, for both *japonica* datasets, derived sSNPs and dSNPs were generally at higher frequencies in putative SS regions, although the effect was not as apparent for sweeps identified with SweeD (Figure S17). Third, like *indica* rice, the SS regions alone did not account for the difference in SFS between *O. rufipogon* and either tropical or temperate *japonica* (*p* ≤ 0.0049 for both comparisons).

### Factors affecting the distribution of variants

Finally, we sought to gain insights into the relative effects of processes that have affected the distribution of genetic variation in Asian rice. To do so, we first devised a measure that is similar to the ‘mean derived allele frequency’ (MDAF) (Simons et al., 2014). Our measure, which we called the ‘mean retained allele frequency’ (MRAF), differs from the MDAF in ignoring the zero class. We ignored the zero class because we were chiefly interested in measuring effects on the set of segregating and fixed variants that were retained through domestication. Similar to the MDAF (Simons et al., 2014), the MRAF was calculated as the average number of derived alleles per individual, divided by twice the number of sites containing derived (segregating sites + fixed) variants within that taxon (Methods). We calculated the MRAF separately for synonymous and putatively deleterious variants. The MRAF was higher for all three rice groups than for the W_15_ *O. rufipogon* sample, regardless of SNP type (Figure 7A).

**Figure 7:**
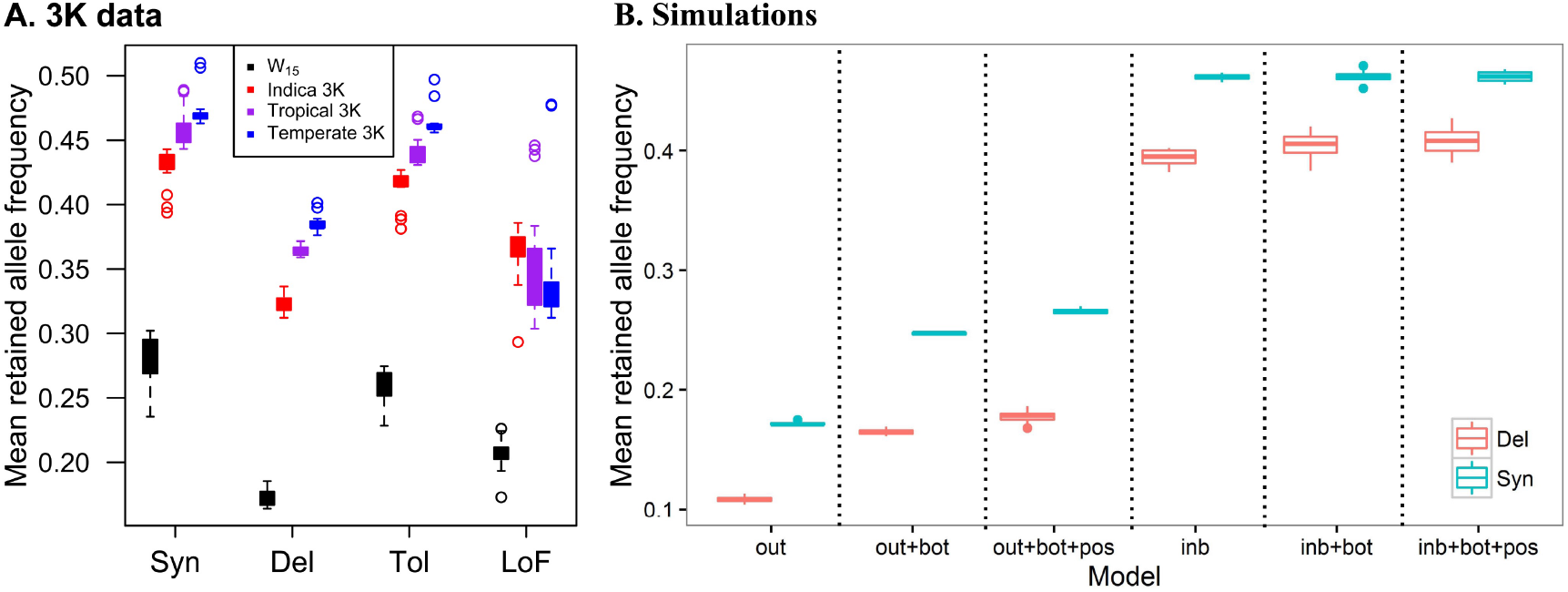
Mean retained allele frequency (MRAF) of empirical (A) and simulated data (B). The x-axis in Fig. 7B defines the six models; ‘`out`’ represents an outbred (random mating) population with a constant population size; ‘`inb`’ represents an inbred (selfing) population with a comparable population size; ‘`bot+out`’ and ‘`bot+inb`’ represents outbred and inbred populations with a domestication bottleneck; ‘`bot+out+pos`’ and ‘`bot+inb+pos`’ include positively selected alleles.

Rice has a complex history that includes a population bottleneck, positive selection and a shift in mating system. We were curious about the relative effects of these evolutionary forces on genetic diversity, as summarized by the MRAF, and so employed forward simulations to model these varied forces. We simulated models with and without a domestication bottleneck, using parameters similar to those inferred from previous study of rice domestication (Caicedo et al., 2007), positive selection, and inbreeding (Methods). To investigate relative effects across different classes of sites, all simulations included both neutral and deleterious variants.

Figure 7B presents simulation results for six models: an outcrossing population (`out`), an outcrossing population with a bottleneck (`out+bot`), an outcrossing population with a bottleneck and positively selected alleles (`out+bot+pos`) and three analogous models that included complete selfing that co-occurs with the bottleneck (`inb, inb+bot, inb+bot+pos`). Focusing first on simulations for outcrossing populations, the MRAF was higher for synonymous compared to deleterious variants, as was found in the empirical data (Figure 7A). The MRAF of both site classes increased under a bottleneck (`out+bot`) and yet again with positive selection (`out+bot+pos`), indicating that both processes drive surviving variants to higher frequency, as expected. Interestingly, as the models progressed from `out` to `out+bot` to `out+bot+pos`, the difference in mean MRAF between synonymous and deleterious variants became larger (from 0.035 to 0.083 to 0.088, respectively).

The inclusion of selfing (`inb`) had a more substantive effect on the shift of the MRAF than the inclusion of either a bottleneck or positive selection (Figure 7B). Under inbreeding models, the inclusion of a population bottleneck (`inb+bot`) and positive selection (`inb+bot+pos`) had no effect on the mean MRAF of synonymous sites (*t*-tests, *p*>0.55). However, the addition of a bottleneck did increase the mean MRAF of deleterious sites (*t*-test, *p*<0.05), such that the difference in mean MRAF between synonymous and deleterious variants became less pronounced from inb (mean difference = 0.067) to `inb+bot` (0.058). In other words, the MRAF of dSNPs was enriched relative to sSNPs as our models progressed from `inb` 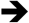 `inb+bot` 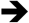 `inb+bot+pos`.

## Discussion

Recent focus on the population genetics of dSNPs in humans (Henn et al., 2015, 2016), plants (Lu et al., 2006, Gunther and Schmid, 2010, Mezmouk and Ross-Ibarra, 2014, Nabholz et al., 2014, Renaut and Rieseberg, 2015, Rodgers-Melnick et al., 2015, Kono et al., 2016) and animals (Schubert et al., 2014, Marsden et al., 2016, Robinson et al., 2016) reflect an emerging recognition that dSNPs may provide unique clues into population history, the dynamics of selection and the genetic bases of phenotypes. This is especially true for the case of domesticated species, where the enrichment of deleterious variants relative to neutral variants reflect a potential “cost of domestication” (Schubert et al., 2014).

Our analyses have provided a snapshot of the fate of deleterious variants during rice domestication. First, dSNPs are typically found at low frequency in wild populations (Figures 1 and 2). Second, many of these low frequency SNPs were lost during domestication, probably due to increased rates of genetic drift during the domestication bottleneck and/or due to inbreeding. The phenomenon of loss is reflected in the large zero class in the SFS of domesticated vs. wild germplasm (Figure 2). Third, the surviving dSNPs shifted toward higher frequency (Figures 1 and 2). Both of these processes – i.e., the loss of rare variants and a shifted SFS – also apply to sSNPs, but our data suggest differential effects on dSNPs vs. sSNPs. This differential effect is evident in the higher proportion of derived dSNPs to sSNPs in domesticated rice than wild rice across most frequency classes (Figure 3), in significant *R_(A/B)_* measures (>1.0) for dSNPs (Figure 3), and in elevated ratios of derived dSNPs/sSNPs per individual (Table 1). For all of these measures, the results were largely consistent between different types of presumably deleterious variants (i.e., dSNPs vs. LoF variants; Figure 3), different methods to predict deleterious SNPs (PROVEAN vs. SIFT; Figures. S4 and S5) and different rice datasets (BH data vs. 3K data).

Our finding that dSNPs are enriched relative to sSNPs is similar to previous observations that have been used to conclude that there is a ‘cost of domestication’ for domesticated crops (Lu et al., 2006; Guther and Schmid, 2010; Renaut and Rieseberg, 2015). However, we also find that the *number* of derived deleterious alleles has increased between wild and crop individuals. Rice individuals in the 3K_subset_ data contain ~200 more deleterious alleles than individuals in the W_15_ *O. rufipogon* sample, a ~3-4% increase (Table 1). To our knowledge, this is the first observation of increased numbers of deleterious variants within domesticated crops, but these results are not dissimilar to breed dogs. Like rice, dog populations contain less genetic diversity than their wolf progenitors, but they also harbor 2.6% more derived deleterious alleles per individual (Marsden et al., 2016). Similarly, serially founded, out-of-Africa human populations exhibit decreasing genetic diversity but increasing counts of derived deleterious variants as a function of geographic distance from Africa (Henn et al., 2016).

### Processes that contribute to enrichment of dSNPs

Several evolutionary forces may contribute to an increase in the number or the proportion of derived deleterious alleles per individual. In out-of-Africa human populations, for example, these factors include range expansion, serial bottlenecks under which moderately deleterious variants evolve as if there were neutral, and differential effects depending on dominance (Henn et al., 2016). At least four major evolutionary factors could drive increased number of deleterious variants in domesticated rice: *i*) changes in population size, particularly bottlenecks associated with domestication (Caicedo et al., 2007, Zhu et al., 2007), *ii*) linked selection (Hartfield and Otto, 2011, Marsden et al., 2016), *iii*) the transition to selfing and *iv*) relaxed selection on wild traits that are no longer important under cultivation (Renaut and Rieseberg, 2015).

Among these, evidence about linkage effects is accumulating. The enrichment of dSNPs in low recombination regions appears to be a general phenomenon, based on studies in *Drosophila* (Campos et al., 2014), humans (Hussin et al., 2015), sunflower (Renaut and Rieseberg, 2015), maize (Rodgers-Melnick et al., 2015), soybean (Kono et al., 2016) and rice (Lu et al., 2006; Figure 4). It remains unclear whether differences between high and low recombination regions of the genome are driven by lower *N_e_* in regions of low recombination (Hill and Robertson, 1966, Felsenstein, 1974a, Charlesworth et al., 1993) or by linkage effects to positively selected variants (Begun and Aquadro, 1992). The relationship between recombination and diversity should be diminished in selfing species (Marais et al., 2004), suggesting that the observed patterns in rice may have accumulated in historically outcrossing *O. rufipogon* populations prior to domestication.

Another aspect of linkage is the enrichment of dSNP frequencies near genes that have experienced selective sweeps (SS). In domesticated dogs, Marsden et al. (2016) document that the average DAC of dSNPs is significantly elevated within SS regions and also that dSNPs experienced the same increase in frequency as sSNPs due to hitchhiking. We find similar effects in rice – i.e., roughly equivalent increases in DACs for dSNPs and sSNPs due to hitchhiking (Figure 6). This suggests that alleles within selected genes, which are presumably of phenotypic importance, may be more often associated with slightly deleterious variants. One must nonetheless be cautious about our approach, because methods that detect SS regions, including π*_d_*/π*_w_*, rely to some extent on a skew of the SFS. This skew should manifest itself as elevated DACs. It is therefore difficult to separate potential methodological artifacts from true signal, but it should be noted that the signal is consistent among SS methods (Figure 6).

Finally, we address the concomitant shift in population size and mating system in rice. It is generally thought that a shift to selfing offers advantages for an incipient crop, such as reproductive assurance, reduced opportunities for gene flow between an incipient crop and its wild ancestor (Dempewolf et al., 2012), and the creation of lines that “breed true” for agronomically advantageous traits (Allard, 1999). This shift may also affect the accumulation of deleterious mutations, but the effect can be difficult to predict, because of antagonistic effects (Arunkumar et al., 2015). On one hand, inbreeding increases homozygosity, exposing recessive deleterious mutations to natural selection (Lande and Schemske, 1985) and potentially leading to the purging of deleterious alleles (Charlesworth and Willis, 2009). On the other hand, inbreeding reduces both population size and effective recombination rates (Nordborg, 2000), thereby reducing the efficiency of selection and contributing to the retention and possible fixation of deleterious variants (Takebayashi and Morrell, 2001).

We have used forward simulations to begin to examine the interplay between inbreeding and demographic (bottleneck) effects under parameters designed to reflect those expected during *O. sativa* domestication. These simulations are unlikely to precisely mimic rice genome history, but they offer insight into the relative effects of evolutionary forces that may have shaped segregating variation in rice. Under outcrossing models, a bottleneck increases the MRAF, as expected, and positive selection increases it even further for both deleterious and synonymous variants (Figure 7B). Under the selfing model, the MRAF of synonymous sites increased dramatically immediately. The addition of a bottleneck and positive selection enriched the MRAF of deleterious variants, but not synonymous variants, such that the MRAF of synonymous and deleterious variants became more similar (Figure 7B).

To the extent that these are representative models, they suggest that the observed difference in MRAFs between *O. rufipogon* and domesticated rice have been affected by selfing more than a bottleneck or positive selection, both of which have subtle effects in the presence of inbreeding (Figure 7B). A relevant comparison is to dog domestication, which occurred in two stages: a population bottleneck associated with domestication ~15,000 years ago (Vonholdt et al., 2010) and inbreeding within the last few hundred years to produce modern breeds. In this case, the domestication bottleneck, rather than inbreeding, has had a larger effect on the accumulation of deleterious genetic variation (Marsden et al, 2016), perhaps because inbreeding in dogs has been more recent and not as intense as inbreeding in rice. Our simulations suggest that inbreeding has had the larger effect in rice, but this is also dependent on assumptions in our models. We have, for example, assumed that selfing was coincident with the domestication bottleneck, but we cannot know this with certainly, especially given the lengthy ‘pre-domestication’ of some crops (Purugganan and Fuller, 2009, Meyer et al., 2016). We have also made assumptions about population sizes, the timing of demographic events, recovery times from those events (Brandvain and Wright, 2016), dominance coefficients (*h*=0.5), and patterns of positive selection. In the future, it will be important to vary these parameter values to better understand their potential effects on crop diversity and the potential cost of domestication.

### Caveats and Assumptions

We close with consideration of the caveats and assumptions of our analyses. While we have tried to avoid potential pitfalls by using multiple approaches (different datasets, SNP calling methods, dSNP predictors, and SS inference metrics), important limitations remain. One is potential reference bias, because the use of the *japonica* reference is expected to decrease the probability that a *japonica* variant (as opposed to an *indica* variant) returns a low PROVEAN or SIFT score (Lohmueller et al., 2008). We have adjusted for this bias by submitting the ancestral allele — rather than the reference allele — to annotation programs (Kono et al., 2016). Without this adjustment, a reference bias was patently obvious, because the SFS of *japonica* dSNPs lacked a high frequency peak, and the U-shape of tSNPs became commensurately more extreme. We cannot know that we have corrected completely for reference bias but do advocate caution when interpreting results from dSNP studies that make no attempt to correct for reference bias. The effect can be substantial.

Our treatment of reference bias requires accurate inference of the ancestral state of variants. To date, most population genetic studies of Asian rice have relied on outgroup sequences from *O. meridionalis* e.g., (Caicedo et al., 2007, Gunther and Schmid, 2010), a species that diverged from *O. sativa* ~2 million years ago (Zhu and Ge, 2005). When we used *O. meridionalis* as the sole outgroup, we inferred a U-shaped SFS in wild *O. rufipogon,* which is suggestive of consistent parsimony misinference of the ancestral state (Keightley et al., 2016). We instead inferred ancestral states relative to a dataset of 93 accessions of African wild rice (O. *barthii*) (Wang et al., 2014). *O. barthii* is closer phylogenetically to *O. sativa* than *O. meridonalis*, but *O. barthii* sequences form clades distinct from *O. sativa* (Zhu and Ge, 2005). Even so, we have found that ~10% of SNPs sites with minor allele frequencies > 5% are shared between African wild rice and Asian rice, perhaps due to introgression (Huang et al., 2015, but see Wang et al. 2014).

We do not believe that the use of *O. barthii* has distorted our primary inferences, for two reasons. First, systematic misinference of the ancestral state should lead to a U-shaped SFS, which is not observed in *O. rufipogon.* Instead, the U-shaped SFS is unique to *O. sativa* and differentiates wild from domesticated species. Second, we have confirmed our inferences by using *O. meridonalis* and *O. barthii* together as outgroups (Keightley et al., 2016), considering only the sites where the two agree on the ancestral state. The use of two outgroups decreases the number of SNPs with ancestral states by ~10% and ~15% for the BH and 3K datasets, but all analyses based on these reduced SNP sets were qualitatively identical to those with only an *O. barthii* outgroup (e.g., Figure S18).

Finally, we focus briefly on the locations of SS regions identified by three different methods (Figure 5 and FiguresS6 to 16), which rarely overlapped (Table 2). In other words, the three methods identified almost completely independent regions of the rice genome. The lack of convergence among methods may reflect that different tests are designed to capture different signals of selection. However, the results are also sobering, because overlaps in SS regions have been used by a number of groups to argue for or against independent domestication of *indica* and *japonica* rice (He et al., 2011, Molina et al., 2011). Recently, both Huang et al. (2012b) and Civian et al. (2015) have argued for independent domestication events for *japonica* and *indica* based on the observation that there is little overlap in SS regions between the two taxa [also see (Huang and Han, 2015).] The fact that we find little overlap among SS regions identified by distinct methods mirrors the lack of overlap of SS regions identified across the human genome by various studies (Akey, 2009), between domesticated grasses (Gaut, 2015), and between independent domestication events of common bean (Gaut, 2015). Because the inferred locations of SS regions vary markedly by method, sampling and taxon, they should be interpreted with caution, particularly as markers of independent domestication events.

## Materials and Methods

### Sequence polymorphism data

All of the data used in this study are publicly available. Illumina paired-end reads for the BH and 3K dataset were downloaded from the European Nucleotide Archive (ENA; http://www.ebi.ac.uk/ena) (see Tables S1 and S2 for accession numbers). The 3K accessions were chosen randomly among the total set of accessions with >12X coverage for an equal representation (*n*=15 for each set) of *indica*, tropical *japonica* and temperate *japonica* rice accessions. We also downloaded resequencing reads from *O. barthii* to polarize SNPs as either ancestral or derived. Sequencing reads for 93 *O. barthii* accessions (Wang et al., 2014) were obtained from the Sequence Read Archive (SRA) database of the National Center for Biotechnology Information (NCBI; http://www.ncbi.nlm.nih.gov/sra/) (see Table S4 for accession numbers). Sequencing reads for another outgroup taxon, *O. meridonalis* were obtained from NCBI (BioProject No: PRJNA264483) (Zhang et al., 2014).

### Read alignment and SNP detection

Paired-end reads for *O. sativa* and *O. rufipogon* data were assessed for quality using FastQC V0.11.2, and then preprocessed to filter adapter contamination and low quality bases using Trimmomatic V0.32 (Bolger et al., 2014). The trimmed reads were mapped to the reference genome for *japonica* Nipponbare rice (MSU V7), which was downloaded from the Rice Genome Annotation Project (http://rice.plantbiology.msu.edu). Mapping was performed with the ALN and SAMPE commands implemented in the software Burrows-Wheeler Aligner (BWA) V0.7.8 (Li and Durbin, 2010), using default parameters. All reads with a mapping quality score of < 30 were discarded.

The method of SNP calling varied with the dataset. For the BH data, alignment files from BWA mapping were processed further by removing PCR duplicates and by conducting indel realignments using Picard tools V1.96 (http://sourceforge.net/projects/picard/files/picard-tools/1.96/) and GATK V3.1 (McKenna et al., 2010), and then used as input for ANGSD V0.901, which is designed to deal with sequences of low depth (Korneliussen et al., 2014). ANGSD was run with the command line:

~~~
angsd -b BAMLIST -anc OUTGROUP –out OUTFILE -remove_bads -uniqueOnly 1
-minMapQ 30 -minQ 20 -only_proper_pairs 1 -trim 0 -minInd NUMBER -P
CPUNUMBERS -setMinDepth 3 -setMaxDepth 15 -GL 1 -doSaf 1 -doMaf 2
-SNP_pval 1e-3 -doMajorMinor 1 -baq 1 –C 50 –ref REFSEQ
~~~

We considered only SNPs that had between 3X and 15X coverage, with the high-end implemented to avoid regions with copy number variation (Huang et al., 2012b). For SNP calling, we used only uniquely mapping reads, and bases with quality score of < 20 were removed. SNP sites with >50% missing data were discarded.

For the higher coverage ‘3K’ dataset, we used SAMtools V1.2 (Li et al., 2009) and GATK V3.1 to call SNPs. After mapping reads of each accession onto the reference genome, alignments were merged and potential PCR duplications were removed using Picard tools V1.96. Unmapped and non-unique reads were filtered using SAMtools V1.2. We realigned reads near indels by using the IndelRealigner and BaseRecalibrator packages in GATK to minimize the number of mismatched bases. The resulting mapping alignments were used as input for UnifiedGenotyper package in GATK and for SAMtools. SNPs that were identified by both tools, with no missing data and a minimum phred-scaled confidence threshold of 50, were retained. Subsequently, SNP calls were further refined by using the VariantRecalibrator and ApplyRecalibration packages in GATK on the basis of two sets of “known” rice SNPs (9,713,967 and 2,593,842) that were downloaded from the dbSNP and SNP-Seek databases (Alexandrov et al., 2015). These same SNP detection methods were applied to the subset of 29 *O. rufipogon*with >4X coverage that were used as the diversity panel to infer SS regions (Table S1), although no prior variants were available.

Finally, sequence reads for the outgroup dataset were aligned to the reference genome using stampy V1.0.21 (Lunter and Goodson, 2011), and then a pseudo-ancestral genome sequence was created using ANGSD (Korneliussen et al., 2014) with the parameters “`-doFasta 2 -doCounts 1`”. This pseudo-ancestral genome was used to determine the ancestral state of each SNP in *O. sativa* and *O. rufipogon*.

### SNP annotation and deleterious mutation prediction

SNPs were annotated using the latest version of ANNOVAR (Wang et al., 2010) relative to the *japonica* reference genome (MSU v 7.0). SNPs were annotated as synonymous, nonsynonymous, intergenic, splicing, stop-gain and stop-loss related. Throughout the study, we combined SNPs that contribute to splicing variation, stop-gain and stop-loss and called them loss-of-function (LoF) mutations.

To discriminate putatively deleterious nSNPs from tolerant nSNPs, nSNPs were predicted as deleterious or tolerated using PROVEAN V1.1.5 against a search of the NCBI nr protein database (Choi et al., 2012). To reduce the effects of reference bias, predictions of deleterious variants were inferred using the ancestral (rather than the reference) variant. Following previous convention (Renaut and Rieseberg, 2015), we considered an nSNP to be a deleterious dSNP if it had a PROVEAN score ≤ −2.5 and a tolerant tSNP when a PROVEAN score was > −2.5. To assess consistency, we also employed SIFT (Kumar et al., 2009) to predict nSNPs as dSNPs or tSNPs. For these analyses, a nSNP was defined as a dSNP if it had a normalized probability < 0.05, and an nSNP was predicted to be a tSNP with a SIFT score ≥ 0.05.

### Calculating site frequency spectra

Following Huang et al. (2012b), we separated the BH dataset of 1,212 accessions into five populations: *indica*, *japonica* (mostly temperate) and three *O. rufipogon* subpopulations (W_I_, W_II_, and W_III_). The five subpopulations were composed of 436, 330, 155, 121, and 170 individuals, respectively (Table S1).

To calculate the site frequency spectrum (SFS) for BH subpopulations, we initially projected the sample size of all five subpopulations to that smallest W_II_ population of *n*=121. However, many of the 121 accessions had low sequencing depth and high levels of missing data. We therefore focused on the W_II_ population to find criteria suitable for inclusion. Ultimately, we sought to retain ≥ 90% of SNP sites within each SNP category, which resulted in a sample size of *n* = 70 for the W_II_ population. Accordingly, we randomly sampled *n* = 70 individuals from the remaining four subpopulations, so long as the sample retained ≥ 90% of SNP sites for each category, to mimic the W_II_ sample.

Given a sample of *n* = 70 for each of the five subpopulations, the SFS for each subpopulation was calculated using the formula proposed by Nielsen et al., (2005), where the *O. barthii* sequence was used as an outgroup to determine the polarity of the mutations.

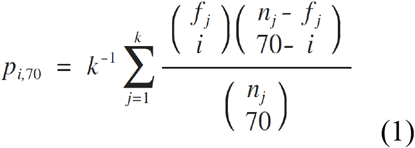

In this formula (1), *p_i,70_*, represents the hypergeometric probability of the derived allele frequency (DAF) of SNPs found in *i* individuals in a sample size of 70; *k* is the total number of SNPs in the dataset; *n_j_* and *f_j_* are the sample size and the number of derived alleles of the *j*th SNP, respectively. The SFS for the 3K data were calculated by focusing on a common set of SNPs that had no missing data and that were segregating in the total population of *n*=60 individuals. The SFS for sSNPs, tSNPs, dSNPs and LoF SNPs were compared with the Kolmogorov-Smirnov test, based on proportions of SNPs at different frequencies.

### R_A/B_ – A relative measure of dSNPs frequency enhancement

We adopted a metric to assess the accumulation of deleterious variants in either cultivated or wild rice populations (Xue et al., 2015). In this analysis, the statistic *L*_A,B_(C) compares two populations (A and B) within a given particular category, *C*, of SNP sites (e.g., dSNPs). It was calculated by counting the derived alleles found at specific sites in population A rather than B and then normalized by the same metric calculated in synonymous sites (S). The calculation of *L*_A,B_(*C*) was:

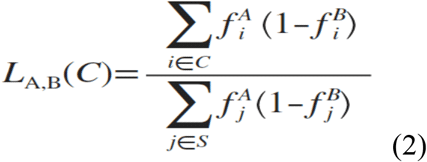

where 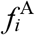 and 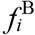 are the observed derived allele frequency at each site *i* in populations A and B, respectively, and *S* refers to sSNPs. The ratio *R*_A/B_(*C*) = *L*_A,B_(*C*) / *L*_B,A_(*C*) then measures the relative number of derived alleles that occur more often in population A than that in population B. To obtain the standard errors of *R*_A/B_(*C*) we used the weighted-block jackknife method (Kunsch, 1989), where each of the tested SNP datasets was divided into 50 contiguous blocks and then the *R*_A/B_(*C*) values were recomputed. A *P* value was assigned by using a *Z* score assuming a normal distribution (Do et al., 2015).

### Calculation of recombination rate

The high-density rice genetic map was downloaded from http://rgp.dna.affrc.go.jp/E/publicdata/geneticmap2000/index.html, on which a total of 3,267 EST markers were anchored. We extracted the sequences of these markers from the dbEST database in NCBI, which were used as query to perform a BLAST search against the rice genome sequences (MSU V7) to annotate their physical positions. Finally, we normalized the recombination rate to centiMorgans (cM) per 100kb between different markers, and then calculated the average recombination rate in 3 or 2MB window segments for the BH and 3K datasets.

### Identification of selective sweep regions

Both SweeD (Pavlidis et al., 2013) and XP-CLR (Chen et al., 2010) were used for identifying selective sweep (SS) regions separately in *indica* and *japonica* populations. SweeD was used with a sliding window size of 10kb, and the *O. barthii* genome sequence (Zhang et al., 2014) was used as an outgroup to determine whether alleles were ancestral or derived. XP-CLR was applied to the 3K datasets along with a subset of 29 *O. rufipogon* individuals that had > 4X coverage and for which we could infer explicit genotypes (Table S1). Both packages were applied with 5% cutoffs to define putative sweep regions. The chromosomal regions identified by SweeD and XP-CLR are provided in Table S9.

We calculated the percentage of genes overlapping between two sets of SS regions, defined as:

~~~
Overlap%= number of genes in common/ ((number of genes in the first set of SS regions + number of genes in the second set of SS regions)-number of genes in common))*100
~~~

### Forward simulations and MRAF

We conducted forward simulations using the software SLiM V1.8 (Messer, 2013). SLiM includes both selection and linkage in a Wright-Fisher model with non-overlapping generations. Similar to previous demographic studies of Asian rice domestication (Caicedo et al., 2007), we simulated a population of *N* = 10,000 individuals, which were run for 10*N* generations to reach equilibrium. We then introduced a domestication bottleneck of size *N*_b_/*N* = 0.01 at generation 10.1 *N* until generation 10.5 *N*, when the population size recovered to size *N* until the end of the simulation at 11.0 *N* generations. For the selfing populations, the population switched from outcrossing to total inbreeding (inbreeding coefficient *F* = 1) at the beginning of the domestication bottleneck.

All simulations assumed a constant mutation rate (*μ* = 6.5 × 10^-9^ substitutions per site per generation) (Gaut et al., 1996) and recombination rate (*ρ* = 4 × 10^-8^ recombinants per generation) (Gaut et al., 2007) across a single chromosome of 100 Mb with alternating 400 bp of noncoding and 200 bp of coding DNA. All noncoding and 75% of coding sequences were selectively neutral (*s* = 0). The remaining 25% of coding sequences were under negative selection under an additive model, with s following a gamma distribution with shape parameter 0.3 and mean −0.05. This DFE was taken from another study of plant mating system (Arunkumar et al., 2015), but we also estimated the DFE of *O. rufipogon* empirically using dfe-alpha-release-2.15 (Eyre-Walker and Keightley, 2009) and the unfolded SFS of the W_15_ sample. The estimated DFE for wild rice was nearly identical to that from Arunkumar et al. (2015), because s had an estimated shape parameter of 0.28 (95% CI: 0.25 to 0.31) and a mean of −0.048 (95% CI: −0.055 to −0.043). Given the similarities between the estimated and assumed DFE, we performed simulations using only the DFE from Arunkumar et al. (2015).

For the inbreeding model without a bottleneck, we followed the method of (Arunkumar et al., 2015) to adjust population size after the outcrossing-selfing transition by calculating the reduction in silent genetic diversity (*θ*_w_ = 4*N*_e_*μ* where *θ*_w_ is genetic diversity, *N*_e_ is effective population size and *μ* is mutation rate). This makes the measures equivalent and the simulations comparable between the inbreeding and outcrossing models that do not include a population bottleneck or positive selection (i.e, out vs. int; Figure 7B).

For the simulations with positive selection, we introduced 20 predetermined mutations with *s* drawn from an exponential distribution of mean 0.05 at the beginning of domestication. For all mutations under positive or negative selection, we assumed a dominance coefficient *h* = 0.5 (i.e., an additive model).

The results for each model were summarized over 20 separate runs of SLiM; the SLiM input is available as Supplementary Text. The MRAF was calculated for simulated data sets and the subset of 3K data as the sum of derived alleles across sites divided by twice the total number of (segregating sites + fixed sites). Note that this definition varies from that of Simons et al (2014) by not including the zero class.

## Acknowledgements

We would like to thank three anonymous reviewers for contributing comments that greatly improved the manuscript. We thank J. Aguirre, T. Kono, D. Seymour, L. Li, and A. Poets for reading earlier versions of the manuscript, and J. Aguirre for assistance with a figure. J. Ross-Ibarra also provided comments and a revised version of XP-CLR. QL is supported by the National Natural Science Foundation of China (grant no. 31471431) and the Training Program for Outstanding Young Talents of Zhejiang A&F University. YZ is supported by the International Postdoctoral Exchange Fellowship Program 2015 awarded by the Office of China Postdoctoral Council. PLM is supported by NSF Plant Genome Program (DBI-1339393). BSG is supported by NSF IOS-1542703 and the Albert and Elaine Borchard Foundation.

